# Photobiomodulation Does Not Influence Maturation and Leads to Mild Improvements in Functional Healing of Mouse Achilles Tendons

**DOI:** 10.1101/735092

**Authors:** Ryan C. Locke, Elisabeth A. Lemmon, Ellen Dudzinski, Sarah C. Kopa, Julianna M. Wayne, Jaclyn M. Soulas, Luis De Taboada, Megan L. Killian

## Abstract

Tendon rupture can occur at any age and is commonly treated non-operatively, yet can result in persisting symptoms. Thus, a need exists to improve non-operative treatments of injured tendons. Photobiomodulation (PBM) therapy has shown promise in the clinic and is hypothesized to stimulate mitochondrial-related metabolism and improve healing. However, the effect of PBM therapy on mitochondrial function during tendon maturation and healing are unknown, and its effect on tendon structure and function remain unclear. In this study, near-infrared light (980:810nm blend, 2.5J/cm^2^) was applied at low (30mW/cm^2^) or high (300mW/cm^2^) irradiance to unilateral Achilles tendons of CD-1 mice during postnatal growth (maturation) as well as adult mice with bilateral Achilles tenotomy (healing). The chronic effect of PBM therapy on tendon structure and function was determined using histology and mechanics, and the acute effect of PBM therapy on mitochondrial-related gene expression was assessed. During maturation and healing, collagen alignment, cell number, and nuclear shape were unaffected by chronic PBM therapy. We found a sex-dependent effect of PBM therapy during healing on mechanical outcomes (e.g., increased stiffness and Young’s modulus for PBM-treated females, and increased strain at ultimate stress for PBM-treated males). Mitochondria-related gene expression was marginally influenced by PBM therapy for both maturation and healing studies. This study was the first to implement PBM therapy during both growth and healing of the murine tendon. PBM therapy resulted in marginal and sex-dependent effects on murine tendon.

## INTRODUCTION

Injury of the Achilles tendon is one of the most common tendon injuries that effects adolescent and adult populations.^1–3^ Achilles rupture and tendinopathy are often debilitating, resulting in limited mobility, impaired joint function, and elevated risk of long-term disability and pain if left untreated.^1,4,5^ Unfortunately, tendon injuries are difficult to treat.^6,7^ Following surgical repair, the risk of re-rupture of the Achilles tendon is greater than 15% and is elevated in patients under the age of 30.^8–10^ Similarly, non-operative therapies such as non-steroidal anti-inflammatory drugs, platelet-rich plasm injections, and physical therapy regimens often leave patients with persisting symptoms, reduced function, and high re-rupture rates.^11,12^ Failure of the Achilles tendon to adequately heal often results from poor cell-mediated reintegration, remodeling, and regeneration during the healing process.^13^ Therefore, developing and improving rehabilitative tools that can regulate cell behavior during tendon healing are needed to accelerate tendon reintegration following injury and reduce the risk of re-rupture following repair.

Photobiomodulation (PBM) therapy is a clinically-available tool for non-invasive treatment of musculoskeletal injuries. PBM therapy can reduce rehabilitation time and improve treatment outcomes when implemented in physical therapy and outpatient clinics.^14–19^ PBM therapy is the application of near-infrared light to increase cellular metabolism, including ATP^20^ and collagen production^21,22^. PBM use in muscle can activate mitochondria,^23–25^ yet it is unclear if the cellular mechanism of PBM is the same during tendon maturation and healing. Increased mitochondrial signaling can influence many downstream targets that control metabolic activity as well as apoptosis, cell proliferation, and collagen synthesis.^26^ In addition, tissue remodeling and pain may be regulated by the metabolic activity of tissues.^27^ While promising results have been shown for PBM treatment of adult tendon injuries in the clinic,^19^ with small animal models,^28–33^ and in tandem with other therapies,^19,34–36^ the effect of PBM therapy on the maturation of tendon has not yet been explored. This is important, because PBM therapy of adolescent tendon injuries may influence the metabolic and remodeling processes of tendon during growth and therefore influence the long-term health of the tendon. Additionally, while PBM therapy is used to reduce pain and increase return-to function for many patients with tendinopathy^16^ and animal models of tendinopathy^28–33^, little work has systematically investigated the therapeutic potential and functional outcomes of PBM therapy on adult tendon healing.^37^ Here, we investigated how PBM therapy modulates mitochondrial metabolism in tendon and determined if and how PBM therapy can regulate the organization, structure, and function of tendon during maturation and healing.

The purpose of this study was to investigate how PBM therapy affects the maturation of tendon *in vivo*, as well as how PBM therapy modulates the structural and biomechanical characteristics of the healing tendon. Specifically, we 1) assessed the effect of low-, high-, and no-irradiance PBM therapy during tendon maturation and 2) evaluated the efficacy of high versus no irradiance of PBM therapy during adult tendon healing. Using a paired study design (Figure 1), we hypothesized that PBM therapy at low or high power would: acutely increase gene expression related to mitochondria metabolism during tendon maturation and healing, not damage the structural and mechanical properties of the maturing tendon, and improve the structural and mechanical properties of the healing tendon.

**Figure 1.**
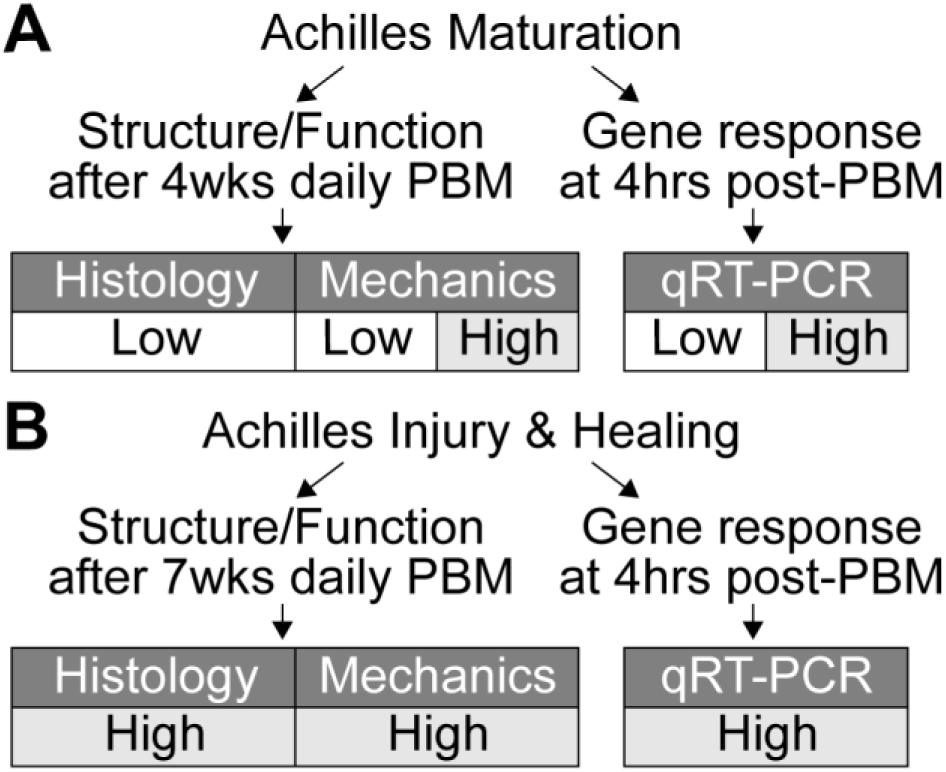
Study design. The effects of photobiomodulation (PBM) on the structure, function, and gene expression of the mouse Achilles tendon during (A) maturation and (B) injury and healing were observed in this study.

## MATERIALS AND METHODS

### Study Design

A total of 54 CD-1 mice (Envigo) were used for all experiments in accordance with approval from the University of Delaware Institutional Animal Care and Use Committee. We designed a two-part study to establish the effect of PBM on Achilles tendon maturation as well as adult healing (Figure 1). Continuous wave of concurrently delivered 980nm and 810nm near-infrared light (80:20 power ratio, 1.5cm^2^ cross sectional area (CSA), hexagon, LightForce FXi Laser, LiteCure) was applied to Achilles tendons unilaterally and the paired contralateral Achilles tendon (untreated) was used as controls for all experiments. For each dose of light, exposure time was controlled to deliver 2.5J/cm^2^ at either 30mW/cm^2^ or 300mW/cm^2^ (Figure 1). The effect of irradiance was tested in the maturation study, and only high irradiance was used for the healing study (Figure 1). The dose parameters were chosen based on previous research.^15^ For all studies, the PBM-dosed tendon (left or right Achilles) was assigned using controlled randomization.

### Bilateral Tenotomy of the Mouse Achilles Tendon

Mice (N = 17) were anesthetized using 3% isoflurane carried by oxygen and buprenorphine (0.01mg/kg) was administered subcutaneously as analgesia. Hair was removed from the skin for subcutaneous exposure of both Achilles tendons using a topical cream. A skin incision was made medially to expose the Achilles tendon. Bilateral Achilles tendons were transected at the midsubstance using #11 scalpel blades. Surgical sites were closed using 5-0 degradable suture (Vicryl, Ethicon Inc., Somerville, New Jersey). Mice were given bupivacaine hydrochloride (0.01mg/kg) for nerve block. Achilles tendons healed for 1-week prior to PBM dosing to allow the sutures to degrade and wound site to close.

### Photobiomodulation Therapy for Mice

Mice were anesthetized via 3% isoflurane carried by oxygen for the duration of PBM dosing, and the anesthesia time was controlled for all experiments (∼10-minutes/dose). Mice were weighed during the dosing periods for the maturation study to make sure that repetitive anesthesia or PBM therapy did not affect their growth (Supplemental Figure 1). Prior to PBM dosing, the hair above the Achilles tendon and calf muscles was removed as needed using a topical cream because hair absorbs a significant amount of light (Supplemental Figure 5).^38^ Mice were then placed in the prone position on a custom-made jig that holds the light probe perpendicular and juxtaposed to the dorsal Achilles tendon in a hands-free manner to reduce variability between doses (Thor Labs, Newton, NJ). Briefly, the dorsal foot was taped down in plantar flexion to keep the Achilles tendon stable during dosing, then the light probe was lowered perpendicular to and directly on top of the dorsal side of the Achilles tendon. The PBM dose was applied, the contralateral limb was manipulated in the same manner but without PBM dosing, and then mice were monitored until awake and alert.

To determine the maturation-dependent effect of PBM irradiance on Achilles tendon structure and function, unilateral Achilles tendons of 2-week old mice (N = 37 total) were dosed with low irradiance (n=13, 7 females and 6 males) or high irradiance (n=8, 4 females and 4 males; functional outcomes only) daily for 4-weeks (Figure 1A). The acute effect of PBM and irradiance during maturation was measured at 4-hours post-treatment of PBM on unilateral Achilles tendons of 2-week old mice that were dosed once with low irradiance (n=6, 3 females and 3 males; n=4, 2 females and 2 males at 24-hours post-PBM) or once with high irradiance (n=6, 3 females and 3 males; Figure 1A and Supplemental Figure 4).

To determine the healing-dependent effect of PBM on Achilles tendons, adult mice (N = 17 total) that underwent bilateral Achilles tenotomy were dosed unilaterally with high irradiance daily for either 7-weeks (structure/function assays; Figure 1B, n=13, 6 females and 7 males) or for 3 days and then sacrificed 4-hours after the third dose (differential gene expression; n=4 females; Figure 1B). Mice were euthanized with carbon dioxide asphyxiation and thoracotomy.

### Tendon Histology

After euthanasia, hindlimbs were scanned using a dental X-ray prior to fixation to compare bone and joint morphology following PBM during maturation (Supplemental Figure 2; Nomad Pro, Aribex). Achilles tendons were dissected and fixed for 24-48hr in 4% paraformaldehyde. Tendons were processed for paraffin sectioning, sectioned in the sagittal plane, then stained using Picrosirius Red (PSR) for collagen organization, 4’,6-diamidino-2-phenylindole (DAPI, NucBlue Fixed, Life Technologies) for cell count and nuclear aspect ratio, and Hematoxylin & Eosin (H&E) for qualitative assessment of tissue morphology (e.g., fibrosis, vessel formation and fat infiltration). Stained sections were imaged using an epi-fluorescent/brightfield microscope (Axio.Observer. Z1, Carl Zeiss, Thornwood, NY). Sections stained with PSR were imaged in the same orientation using circular polarized light microscopy, and the alignment of collagen fibrils was semi-quantitatively scored by four blinded individuals (JS) using the following scoring rubric: 5 = extremely organized and yellow, 4 = predominantly organized and orange, 3 = mixed organization and green, 2 = predominantly random and red/green, 1 = extremely random and red/black. Sections stained with DAPI were imaged at six, non-overlapping locations (0.011mm^2^/image, 40x) along the length of the tendon that served as technical replicates. Cell count and nuclear aspect ratio were quantified by a custom MATLAB code for each technical replicate and then averaged per sample (R2016b, MathWorks, Natick, MA, USA).

**Figure 2.**
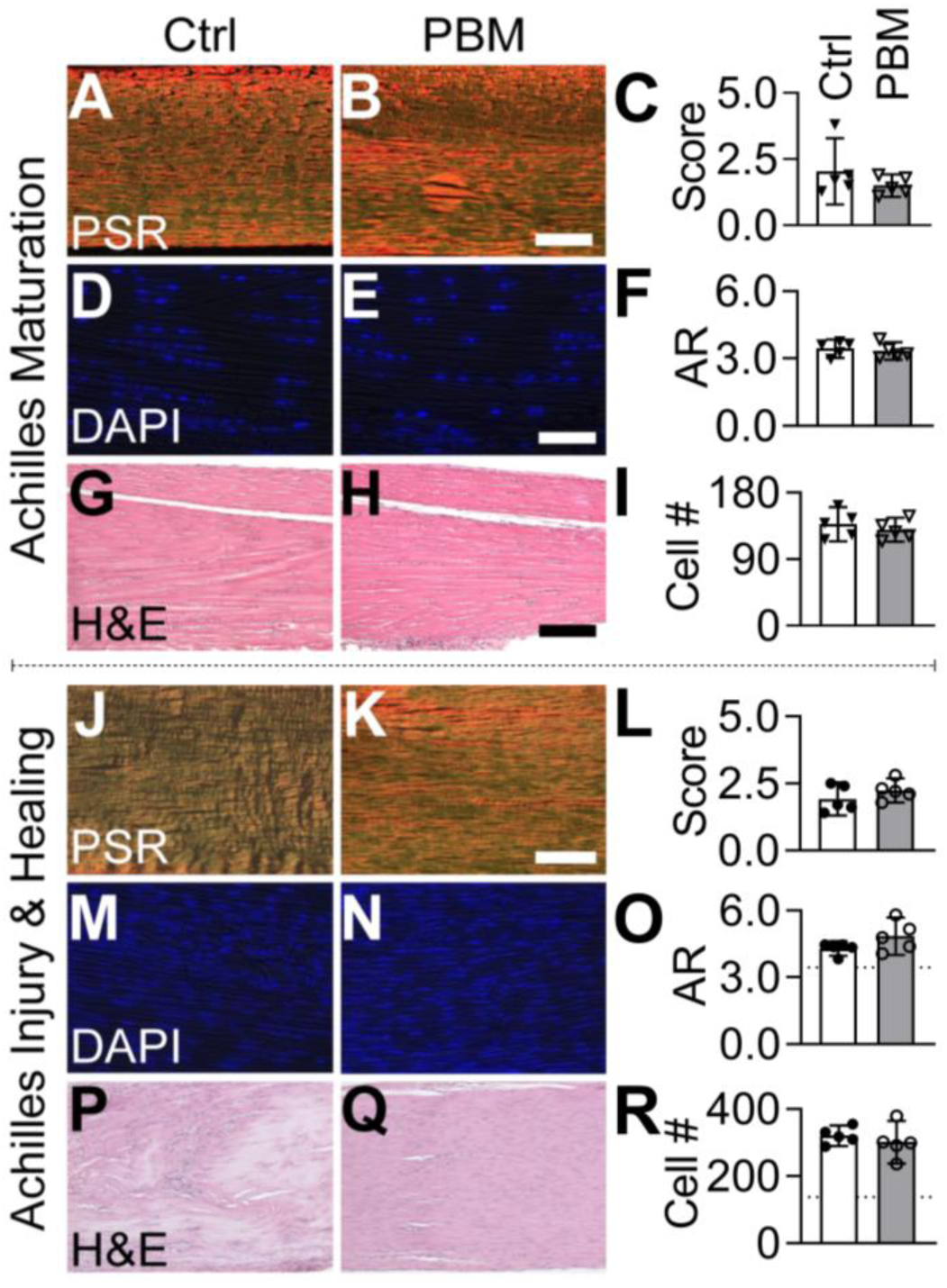
During maturation and adult injury and healing, daily doses of PBM therapy did not affect the structural or cellular properties of the mouse Achilles tendon. (A-C, maturation; J-L, healing) Collagen alignment, (D-F, maturation; M-O, healing) nuclear aspect ratio and (I, maturation; R, healing) cell number, and (G-H, maturation; P-Q healing) fibrosis were observed using Picrosirius red (PSR), DAPI, and hematoxylin and eosin (H&E) stained sections, respectively. Dashed line: Mean of maturation controls (Ctrl). Scale bar for panels B and H: 75µm, Scale bar for panel E and K: 40µm. Data are presented as mean ± standard deviation.

### Tendon Uniaxial Tensile Tests

After euthanasia, mice for biomechanical testing were immediately frozen at -20°C. Prior to mechanical tests, mice were thawed and Achilles tendons were dissected from the proximal and distal tibias. Achilles-calcaneus attachments were left intact, and the gastrocnemius muscle was carefully removed using a #11 scalpel blade. Uniaxial tensile tests were performed (Instron 5943, Norwood, MA) in a PBS bath at room temperature using custom fixtures to ensure uniaxial loading. Mechanical tests included a 0.01N preload, 10 preconditioning cycles from 0.02N to 0.04N at 0.02mm/sec, and ramp to failure at 0.02mm/sec. Cross-sectional areas (CSA) and grip-to-grip gauge lengths were measured after the preload, and CSA were assumed to be ellipsoidal. Load and displacement data were recorded (1kHz acquisition rate), and data were converted to stress and strain based on the CSA and gauge length. Structural and mechanical properties (stiffness, ultimate load, Young’s modulus, ultimate stress, strain at ultimate stress, and area under the curve at ultimate stress [AUC]) were calculated using a custom MATLAB code. Stiffness and Young’s modulus were calculated as the slope of the linear region from the load/displacement and stress/strain curves, respectively. Ultimate load, ultimate stress, and strain at ultimate stress were calculated as the maximum load/stress/strain from their respective curves. AUC was calculated by taking the integral of the stress/strain curve to the ultimate stress.

### Mitochondria Gene Expression in Tendon

To investigate the cellular mechanism of PBM, Achilles tendons were dissected following euthanasia (4-hours after last PBM dose) in RNase-free conditions. Tendons were frozen in liquid nitrogen and stored at -80C. Tissues were pulverized for 30 sec in 2mL tubes with steel balls in liquid nitrogen-cooled blocks (MM400, Retsch, Verder Scientific). RNA was extracted using a commercially available kit (PureLink RNA Mini Kit, Invitrogen). Total RNA was reverse-transcribed using Superscript III VILO (Invitrogen). Quantitative real-time-PCR was performed using RT^2^ Profiler PCR Array for Mouse Mitochondria (330231, Qiagen) with PowerUp SYBR Green (Applied Biosystems) on a LightCycler 96 System (Roche). ΔCT (CT_target gene_ – CT_reference gene_) values were calculated for each gene using *Gusb* as the reference gene (most stable reference gene across all conditions with 30.9±1.1 for maturation and 26.5±0.4 for healing Ct values). Reference gene was selected as the gene that changed the least due to PBM treatment from a panel of 5 reference gene (*Actb, B2m, Gapdh, Gusb*, and *Hsp90ab1*). ΔCT values for each target gene were transformed to 2^-ΔCT^ and then averaged for control and PBM-treated tendons. Fold change was calculated by dividing the average 2^-ΔCT^ of PBM-treated by the average 2^-ΔCT^ of control for each gene.

### Statistical Analysis

All statistical comparisons were made using Prism (v8.2.0, Graphpad, LaJolla, CA). Paired t-test were used to compare the ΔCT of each gene when comparing tendons treated with PBM during maturation (low), maturation (high), and healing to their paired control tendons. For mechanical outcomes, repeated-measure (control vs. PBM tendon) two-way ANOVAs with Sidak’s correction for multiple comparisons (sex differences) were used. Wilcoxon matched-pairs tests were used to compare semi-quantitative scores of collagen alignment. Paired t-tests were used to compare quantitative measures of cell number and nuclear aspect ratio. The fold change and p-value for each gene were log_2_ and -log_10_ transformed, respectively, to visualize results in volcano plots. For genes that were statistically different between control and PBM, the mean ± standard deviation was plotted as log_2_(2^-ΔCT^). Differential gene expression with a fold change less than -1.5 or greater than 1.5 were plotted for paired comparisons. Mechanical properties, collagen alignment score, cell number, and nuclear aspect ratio were plotted as mean ± standard deviation.

## RESULTS

### Daily PBM Does Not Affect Tendon Structure During Postnatal Growth or Healing

During maturation, daily doses of PBM therapy did not affect the structural or cellular morphology of the growing mouse Achilles tendon (Figure 2). Specifically, collagen alignment was not affected by daily PBM during maturation (Figure 2A-C). Additionally, there were no differences in the collagen of transected tendons between PBM treated and untreated tendons following 8-weeks of healing (Figure 2J-L), although during healing the alignment score trended higher in the PBM group compared to paired controls (Figure 2L). Nuclear shape and cell number were not affected by PBM during maturation (Figure 2D-F) and healing (Figure 2M-O). The number of cells and aspect ratio were significantly increased during healing compared to maturation controls (Figure 2O). From H&E stained sections, no differences were observed between control and PBM groups for maturation (Figure 2G-H) and healing, with fibrosis, characterized by fat infiltration and vessel formation, observed in both groups during healing (Figure 2P-R). Daily PBM and repeated anesthesia needed for each dose of PBM therapy did not affect the gross morphology or mouse growth, respectively (Supplemental Figures 1 and 2).

### Daily PBM Does Not Influence the Mechanical Properties of Tendon Maturation and Improves Tendon Healing in a Sex-Dependent Manner

During maturation, daily doses of PBM therapy at both low and high irradiance for 4-weeks had no significant effect on the mechanical or structural properties of the mouse Achilles tendon (Supplemental Figures 3). No significant differences were observed between the low and high PBM groups or compared to their paired controls (Supplemental Figure 3). Additionally, no sex-dependent significant differences were observed during maturation (Supplemental Figures 3).

Contrastingly, during healing, daily doses of PBM therapy for 7-weeks had sex-dependent effects on the mechanical and structural properties of mouse Achilles tendons (Figure 3). Achilles tendons were stiffer for females in the PBM group compared to sex-matched untreated controls (Figure 3A), and Young’s modulus was significantly higher in the female PBM group compared the male PBM group (Figure 3B). Ultimate strain in the male PBM group was significantly higher compared to the male control group and the female PBM group (Figure 3C). No differences were observed for ultimate load, ultimate stress, and AUC at ultimate stress between paired control and PBM-treated tendons. There were no significant differences in mechanical properties when males and females were analyzed together.

**Figure 3.**
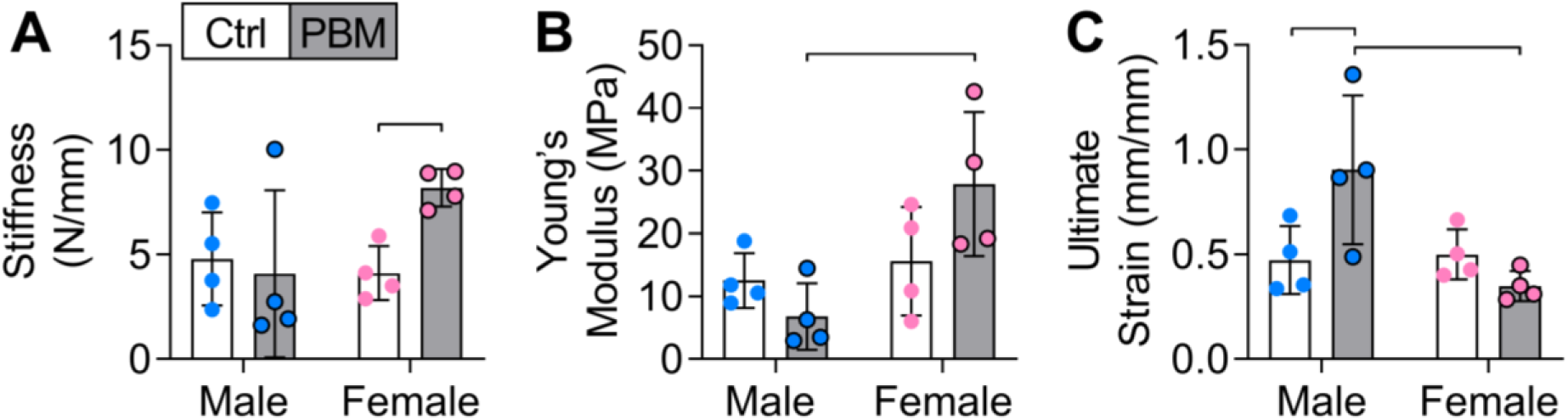
During healing, daily doses of PBM therapy for 7-weeks had sex-dependent effects on the mechanical properties of the mouse Achilles tendon. (A) Stiffness, (B) Young’s modulus, and (C) Ultimate strain for control (Ctrl) and PBM treated tendons of males and females. Data are presented as mean ± standard deviation (p<0.05).

### Acute PBM Marginally Alters Mitochondrial Gene Expression

During both maturation and healing, several genes related to mitochondrial signaling were differentially expressed following PBM treatment (Table 1). During maturation, a single dose of low irradiance PBM treatment downregulated *Bbc3* in females and upregulated *Slc25a23* and *Slc25a24* in males compared to paired controls (Figure 4A). During maturation, a single dose of high irradiance PBM in male tendons led to downregulation of 3 mitochondria-related genes (*Ucp2, Slc25a23*, and *Slc25a13*) compared to paired controls, and this effect was sex-specific (Figure 4B). In females during maturation, high irradiance PBM treatment led to significant upregulation of *Slc25a24* compared to paired controls (Figure 4B). At 24-hours after a single dose of low irradiance PBM, only Slc*25a21* was downregulated by PBM compared to paired controls during maturation (Supplemental Figure 4). During maturation, there were no differences between 4-hours and 24-hours after a single dose of low irradiance PBM, and *Slc25a13* was upregulated in the low group compared to the high group at 4-hours (Supplemental Figure 4). During healing, 3 consecutive days of high irradiance PBM statistically resulted in the significant upregulation of three genes (*Bcl2, Slc25a19*, and *Slc25a37)* and downregulation of *Slc25a13* compared to paired controls (Figure 4C).

**Figure 4.**
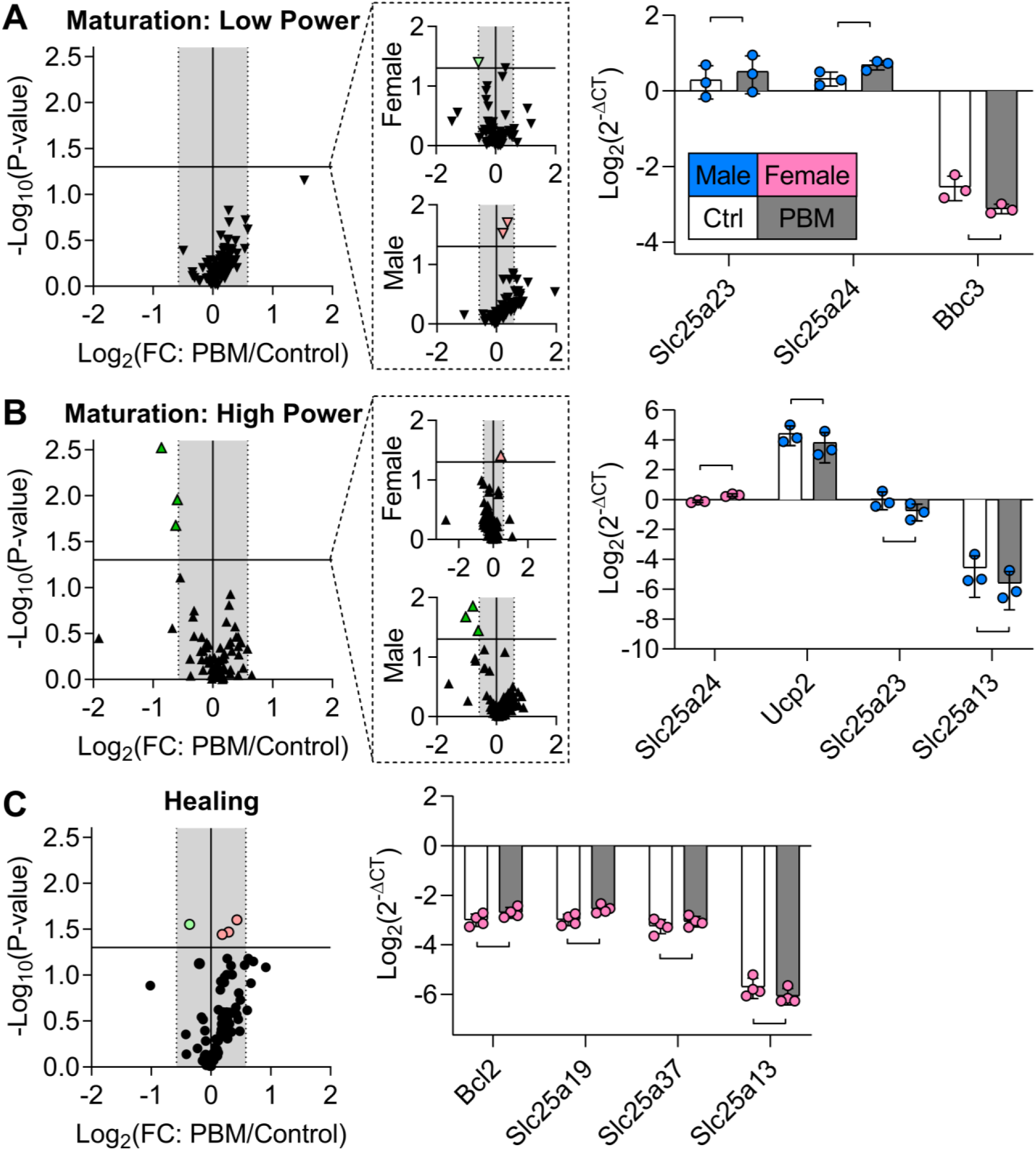
During maturation and healing, mitochondria-related genes were marginally effected by PBM. (A-C) Volcano plots of the acute response from mitochondria-related genes after PBM during maturation (A, low and B, high) and healing (C). Horizontal line: p=0.05, gray box: -1.5 to 1.5-fold change. (A) During maturation, low irradiance PBM significantly downregulated *Bbc3* in females and upregulated *Slc25a23* and *Slc25a24* in males. (B) During maturation, high irradiance PBM downregulated 3 mitochondria-related genes (*Ucp2, Slc25a23*, and *Slc25a13*), and this effect was due to males, not females; only *Slc25a24* was significantly upregulated in females. (C) During healing, high irradiance PBM of females statistically upregulated 3 genes (*Bcl2, Slc25a19*, and *Slc25a37)* and downregulated 1 gene (*Slc25a13*). Data are presented as mean ± standard deviation (p<0.05).

**Table 1.** Genes affected by PBM during tendon maturation and healing (Descriptions from UniProt^42^).

| Sex | Gene | p-value | PBM/Ctrl | Function |
| --- | --- | --- | --- | --- |
| **Maturation: Low Power** |  |  |  |  |
| M | *Slc25a23* | 0.034 | 1.16 | Mediates the transport of Mg-ATP in exchange for phosphate, catalyzing the net uptake or efflux of adenine into or from the inner membrane. Regulates mitochondrial calcium uptake via interaction with MCU and MICU1. |
| M | *Slc25a24* | 0.018 | 1.29 | May play a role in protecting cells against oxidative stress-induced cell death, probably by promoting the formation of calcium-phosphate precipitates in the mitochondrial matrix, and thereby buffering calcium levels in the matrix. |
| F | *Bbc3* | 0.036 | 0.67 | Essential mediator of p53/TP53-dependent and p53/TP53-independent apoptosis. Functions by promoting partial unfolding of BCL2L1 and dissociation of BCL2L1 from p53/TP53. Regulates ER stress-induced neuronal apoptosis. |
| **Maturation: High Power** |  |  |  |  |
| F | *Slc25a24* | 0.039 | 1.339 | Described above. |
| M | *Ucp2* | 0.036 | 0.653 | Mitochondrial transporter protein that creates proton leaks across the inner mitochondrial membrane, thus uncoupling oxidative phosphorylation from ATP synthesis. |
| M | *Slc25a23* | 0.014 | 0.578 | Described above. |
| M | *Slc25a13* | 0.021 | 0.489 | Catalyzes the calcium-dependent exchange of cytoplasmic glutamate with mitochondrial aspartate across the inner membrane. May have a function in the urea cycle. |
| **Healing** |  |  |  |  |
| F | *Bcl2* | 0.034 | 1.228 | Suppresses apoptosis in a variety of cell systems by controlling the mitochondrial membrane permeability. Inhibits caspase activity either by preventing the release of cytochrome c from the mitochondria and/or by binding to the apoptosis-activating factor (APAF1). |
| F | *Slc25a19* | 0.025 | 1.351 | Mitochondrial transporter mediating uptake of thiamine pyrophosphate (ThPP) into mitochondria. |
| F | *Slc25a37* | 0.036 | 1.14 | Mitochondrial iron transporter that plays an essential role in heme biosynthesis. The iron delivered into the mitochondria may be delivered to ferrochelatase to catalyze Fe<sup>2+</sup> incorporation into protoporphyrin IX to make heme. |
| F | *Slc25a13* | 0.028 | 0.78 | Described above. |

## DISCUSSION

This study investigated the role of PBM therapy on the structure and function of the murine Achilles tendon, as well as mechanistically identified that PBM therapy marginally affects mitochondria gene expression, during maturation and healing. We have demonstrated that PBM therapy during tendon maturation does not alter the structural and mechanical properties, regardless of the power level, suggesting that it is a safe therapy for growing tendon. We showed that daily PBM therapy during tendon healing has sex-dependent effects in the mechanical properties of tendon. Mechanistically, we found that PBM therapy marginally influences gene expression related to mitochondrial metabolism.

The mechanism of action of PBM therapy on cells remains unclear and may not be ubiquitous in all cell types. We hypothesized that the initial response of tendon cells to PBM therapy would lead to elevated mitochondria metabolism through the acceptance of photons via photo-acceptors like cytochrome c oxidase (COX).^15,39^ In turn, the accepted photons would lead to increased availability of electrons for the reduction of oxygen, thereby increasing the rate of oxidative phosphorylation and adenosine triphosphate (ATP) synthesis.^15,39^ Our gene expression experiments did not support this hypothesis.

Previous research has shown that PBM therapy can increase cell proliferation, inflammation, extracellular matrix (ECM) remodeling, and collagen synthesis^28–33^. However, ours is the first work to systematically investigate the role of PBM therapy on tendon structure, function, and mitochondrial activity. In the present study, we did not find an overwhelming effect of PBM therapy, with the parameters and times that we used, on tendon-specific mitochondrial metabolism during growth or following injury. These findings may suggest that tendon-specific mitochondria are not sufficiently influenced by PBM therapy. Alternatively, it is possible that the parameters we used did not alter mitochondria metabolism. It is possible that other parameters and altered dosing of PBM therapy may lead to altered mitochondrial-related gene expression.^40^ At the parameters and time points that we assessed, we did identify marginal, statistically significant differences in expression of several mitochondria-related genes induced by PBM therapy during tendon maturation and healing (Table 1).^41^ The majority of differentially-expressed genes that were regulated by PBM therapy were calcium-dependent small molecule transporters that shuttle molecules into mitochondria through the inner membrane, which suggests that calcium signaling may be affected by PBM therapy.^42,43^

During maturation, the sex-dependent effects of PBM therapy (i.e., differential gene expression in males compared to females) are important to consider when establishing the mechanism of action of PBM therapy or other treatment modalities that target mitochondria. Both low and high irradiance PBM therapy regulated *Slc25a23* and *Slc25a24*, and these genes may be targets of PBM therapy in this scenario of tendon maturation.^42,43^ The differences we found between irradiance groups (low versus high) suggest that the irradiance used may alter the mechanism of action of PBM therapy. For example, we found that low irradiance led to upregulated *Slc25a23* and *Slc25a24* and downregulated *Bbc3*, whereas high irradiance led to downregulated *Ucp2, Slc25a23*, and *Slc25a13* and upregulated *Slc25a24*. Although we identified similar genes that were differentially expressed due to low and high irradiance PBM therapy, these changes did not lead to changes in the structure or mechanical properties of the tendon during maturation, as hypothesized.

During healing, we observed increased mechanical properties as well as differentially expressed genes in females, but we did not observe changes in the tendon structure. The genes that we found to be differentially expressed following PBM therapy in the healing context may have enhanced the functional remodeling of the ruptured tendon. We found that, in females, high irradiance PBM therapy during both maturation and tendon healing led to downregulated *Slc25a13. Slc25a13* encodes instructions to make the protein CITRIN, which transports cytoplasmic glutamate for mitochondrial aspartate across the inner membrane, a key feature of the citric acid cycle. Downregulation of *Slc25a13* indicates that the citric acid cycle would rely more on pyruvate than glutamate for production of amino acids and ATP, but it is unclear how this would benefit the healing of tendon. Additionally, during healing, PBM therapy increased expression of *Bcl2*, which is known to inhibit apoptosis by regulating cytochrome c release from mitochondria or by interacting with *Apaf1*.^44^ PBM therapy during healing also upregulated *Slc25a19* and *Slc25a37*, which respectively function to transport thiamine pyrophosphate (ThPP)^45^, a critical coenzyme in the citric acid cycle, and iron, a necessity for the synthesis of heme^46^, into mitochondria through the inner membrane. Yet, it is unclear if the differential expression of these genes led to changes in mechanical outcomes (e.g., stiffness) of healing tendons.

### Clinical implications

In the clinic, PBM therapy alone or in conjunction with traditional interventions, such as physical therapy, has shown promising results for the treatment of tendon injuries in both human and veterinary medicine.^19,47^ Our results corroborate that PBM therapy may be beneficial for tendon healing, and that PBM therapy does not alter Achilles tendon maturation. Our findings support that PBM therapy is a safe treatment modality for growing tissues and that PBM may improve the mechanical properties of the healing adult tendon. Recently, PBM therapy was shown to not alter the morphology or mechanics of human tendon^48^, and our results support these findings. Several studies in humans have shown that repetitive PBM therapy improves the healing process, function, and pain levels of injured tendons.^16,19,27^ These studies highlight the importance of using defined parameters and dosing that warrant beneficial effects on healing. PBM therapy in the clinic and in our study may be beneficial for tendon healing because the functional remodeling of tendon during healing improves without adverse effects.

### Limitations

The effect of PBM therapy on gene expression may be dependent on the energy density applied, yet we only used one energy density for all studies thus other energy densities may have a greater effect on gene expression. Additionally, during maturation, more doses may have amplified or altered gene expression further, yet we quantified the effect of repeated doses in the healing study and did not find an amplified effect. Further, fewer doses might have altered gene expression differently during healing. Gene expression is highly time-dependent, thus we also analyzed gene expression at 24-hours after PBM therapy but did not find meaningful differences compared to 4-hours. A second limitation to this study is that we did not identify structural changes in tendon that may have led to improvements in mechanical properties. Future work may use more quantitative measures of tendon structure, such as second harmonic generation imaging, as well as study the direct role of mitochondria in the healing response of tendon.

## CONCLUSION

This was the first study to systematically assess the paired effect of PBM therapy on the cellular, structural, and mechanical properties of tendon during maturation and healing. This study shows that the mechanism of action of PBM therapy may not be via mitochondria, that PBM therapy does not damage the maturation of tendon, and that PBM therapy mildly improves the healing of ruptured tendon in sex-dependent manner.

## ACKNOWLEDGMENTS

Delaware Bioscience Center for Advanced Technology; LiteCure donation of IR laser; Elahe Ganji for preliminary help with Achilles tensile tests.

