## Supplemental Data for "Photobiomodulation Does Not Influence Maturation and Leads to Mild Improvements in Functional Healing of Mouse Achilles Tendons"

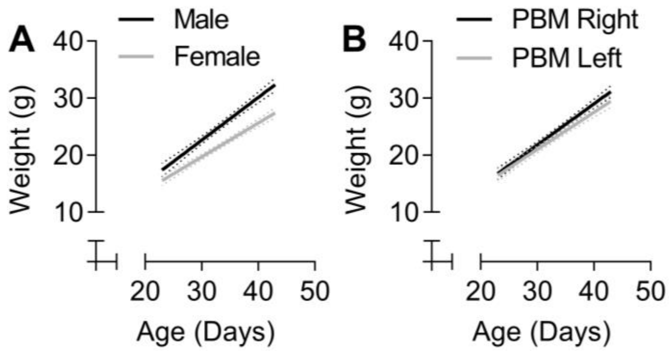


**Supplemental Figure 1. During maturation with repeated anesthesia, male and female mice continued to gain weight.** (A) Males were significantly heavier than females, and (B) no difference was observed for animals treated on the right leg or the left leg. PBM did not negligibly effect the growth of the mice. Dotted line: 95% confidence interval.

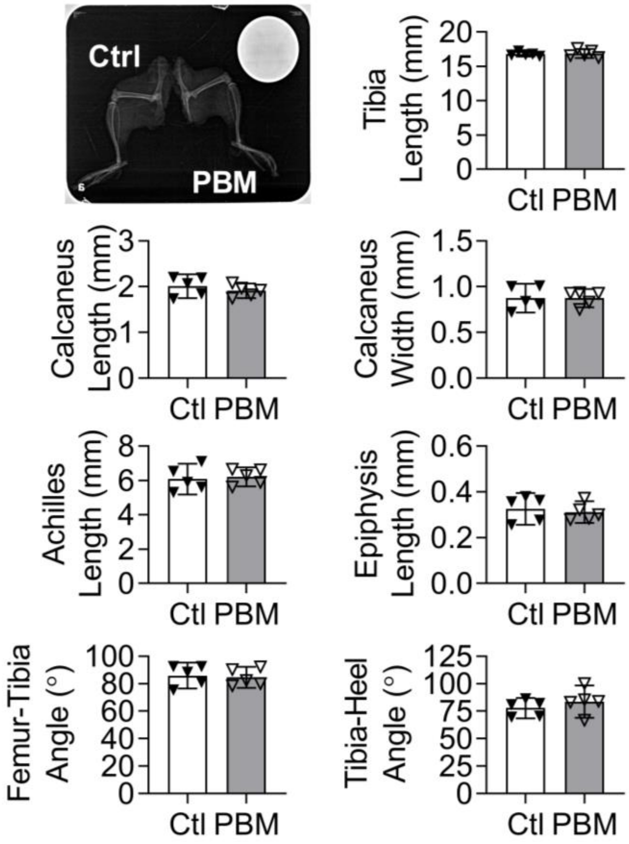


**Supplemental Figure 2. During maturation, low power PBM does not alter joint or bony geometry.** Data are presented as mean irradiance ± standard deviation ($p<0.05$).

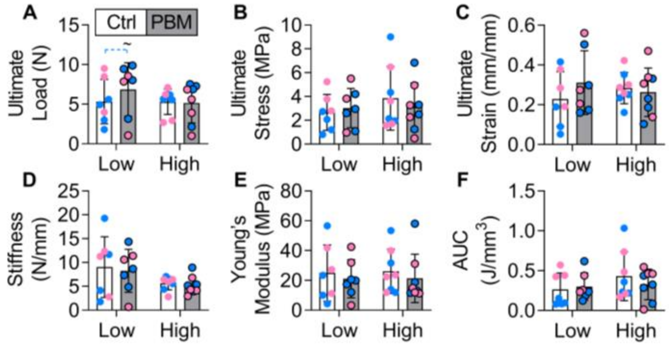


**Supplemental Figure 3. During maturation, daily doses of PBM therapy (low or high irradiance) for 4-weeks did not affect the mechanical and structural properties of the mouse Achilles, regardless of sex.** A) However, a trend of increased ultimate load in males treated with PBM compared to their paired controls was observed (p=0.08) and compared to females treated with PBM (p=0.07). Data are presented as mean ± standard deviation (p<0.05).

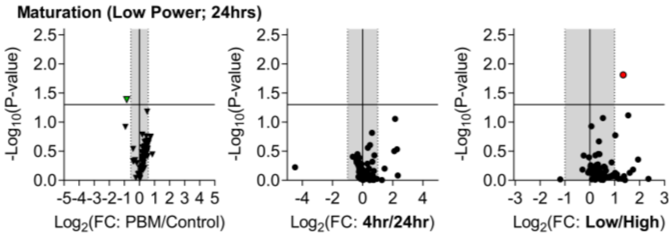


**Supplemental Figure 4. During maturation, negligible differences in gene expression were observed between 4-hours and 24-hours and between low and high.** *Slc25a21* (green point) and *Slc25a13* (red point).

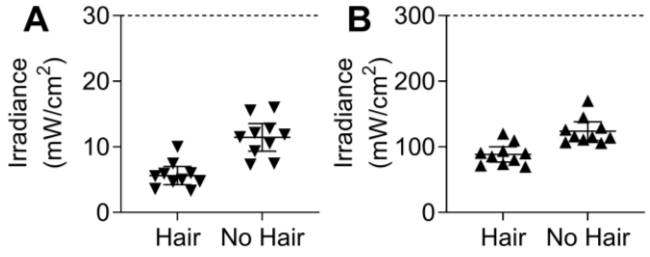


**Supplemental Figure 5. Skin and hair reduces the amount of irradiance that penetrates to the target tissue.** Dotted line: applied irradiance. Data are presented as mean irradiance ± standard deviation (p<0.05).
